# BaCoN (Balanced Correlation Network) improves prediction of gene buffering

**DOI:** 10.1101/2024.07.01.601598

**Authors:** Thomas Rohde, Talip Yasir Demirtas, Angela Helen Shaw, Maximilian Billmann

## Abstract

Buffering between genes is fundamental for robust cellular functions. While experimentally testing all possible gene pairs is infeasible, gene buffering can be predicted genome-wide under the assumption that a gene’s buffering capacity depends on its expression level and the absence of this buffering capacity primes a severe fitness phenotype of the buffered gene. We developed BaCoN (<u>Ba</u>lanced <u>Co</u>rrelation <u>N</u>etwork), a post-hoc unsupervised correction method that amplifies specific signals in expression-vs-fitness effect correlation-based networks. We quantified 147 million potential buffering relationships by associating CRISPR-Cas9-screening fitness effects with transcriptomic data across 1019 Cancer Dependency Map (DepMap) cell lines. BaCoN outperformed state-of-the-art methods including multiple linear regression, based on our newly compiled metrics for gene buffering predictions. Combining BaCoN with batch correction or Cholesky data whitening further boosts predictive performance. We characterized a high-confidence list of 899 buffering predictions and found that while buffering genes overall are often syntenic, buffering paralogs are on different chromosomes. BaCoN performance increases with more screens and genes considered, making it a valuable tool for gene buffering predictions from the constantly growing DepMap.

## Introduction

To ensure robustness of cellular functions, organisms across all phyla rely on genetic redundancy where one gene can compensate for another gene’s loss of function. This genetic redundancy is more complex in higher organisms and genome duplication throughout evolution coupled with retention of genes if beneficial, gave rise to a large number of paralogs (Singh & Isambert, 2019). While the concept of genome duplication, retention of genes and subsequent sequence divergence is an attractive model to predict functional redundancy leading to functional buffering (Ryan *et al*, 2023b; De Kegel & Ryan, 2023), recent studies have shown that, while strongly enriched, the vast majority of paralogs do not functional buffer each other (Esmaeili Anvar *et al*, 2024). At the same time, functional buffering in a wider sense can also be observed between non-paralogs (Ryan *et al*, 2023a; Kryukov *et al*, 2016; Behan *et al*, 2019). For instance, a cell can to cope with either loss of *BRCA1* or *PARP1*, but the combinatorial perturbation of the two genes is synthetic lethal (SL), resembling one of the most well-known clinical implications of functional buffering where *PAPR1* inhibition can help patient harboring *BRCA1* mutations (Farmer *et al*, 2005). While isogenic CRISPR screens allow for testing the genome against one specific mutation, the all-by-all gene search space that cannot be experimentally investigated (Oser *et al*, 2019; Williamson *et al*, 2016; Aregger *et al*, 2020; DeWeirdt *et al*, 2021).

Genetic networks from functional genomics data can assign function to genes in a systematic fashion. Similarity between expression (co-expression) or genetic interaction signatures have been exploited to classify genes by their function in model organisms (Costanzo *et al*, 2016; Fischer *et al*, 2015; Collins *et al*, 2007; Eisen *et al*, 1998; Billmann *et al*, 2018; Costanzo *et al*, 2010). Following this concept, co-essentiality networks in human cells derived from genome-scale CRISPR screens from the Cancer Dependency Map (DepMap) enable systematic prediction of gene function as well (Wainberg *et al*, 2021; Hassan *et al*, 2023; Pan *et al*, 2022). The DepMap perturbs all approximately 18,000 genes in the human genome followed by the measurement of the effect of the perturbation on cell fitness. To date, the DepMap has screened more than 1000 cultured human cell lines and aims to expand this effort to several thousand cell models (Boehm *et al*, 2021). Experimental efforts are accompanied by a continued improvement of the analytical pipeline (Meyers *et al*, 2017; Dempster *et al*, 2021; Iorio *et al*, 2018), which adjusts the data for various factors that would otherwise compromise the interpretation of per-cell line fitness effects as well as co-essentiality networks that measure similarity of fitness effects between genes across all cell lines. A prominent artifact is copy number-driven DNA cutting toxicity effects that confound fitness scores and create a syntenic gene clustering bias (Wainberg *et al*, 2021; Meyers *et al*, 2017). Those analytical pipelines have been instrumental for extracting gene fitness effects from every screened cell line.

A separate class of data normalization methods exploited the CRISPR screening-derived gene fitness effects to generate co-essentiality networks that predict similarity of gene functions (Hassan *et al*, 2023; Wainberg *et al*, 2021; Rahman *et al*, 2021; Boyle *et al*, 2018). A comparison of different normalization strategies showed that the perhaps most crucial part of the normalization strategy is the whitening of the fitness effect gene-by-cell line matrix prior to the computation of between-gene association indices (Gheorghe & Hart, 2022).

More recently, to predict gene buffering, DepMap gene fitness effects have been contrasted with the mRNA expression in matched panels of cell lines. Conceptually, such a contrast would show positive association indices if the potentially buffered gene is more essential in cell lines in which the buffering gene is less expressed (De Kegel *et al*, 2021; Köferle *et al*, 2022; Pacini *et al*, 2024; Krieg *et al*, 2024; Lord *et al*, 2020) (Figure 1a). In this case, gene fitness effects and mRNA expression must be measured in the same cell lines but in independent experiments. Multiple linear regression models are routinely deployed to estimate association indices while accounting for confounding factors such as cell lineage or growth media conditions (Pacini *et al*, 2024; De Kegel *et al*, 2021). Other powerful methods that perform data normalization prior to computing the association indices, such as data whitening, may carry the risk of unlinking biologically relevant co-variation between the data sets. The benefit and risk of such a priori data normalization methods when predicting gene buffering from the DepMap have not been explored.

**Figure 1.**
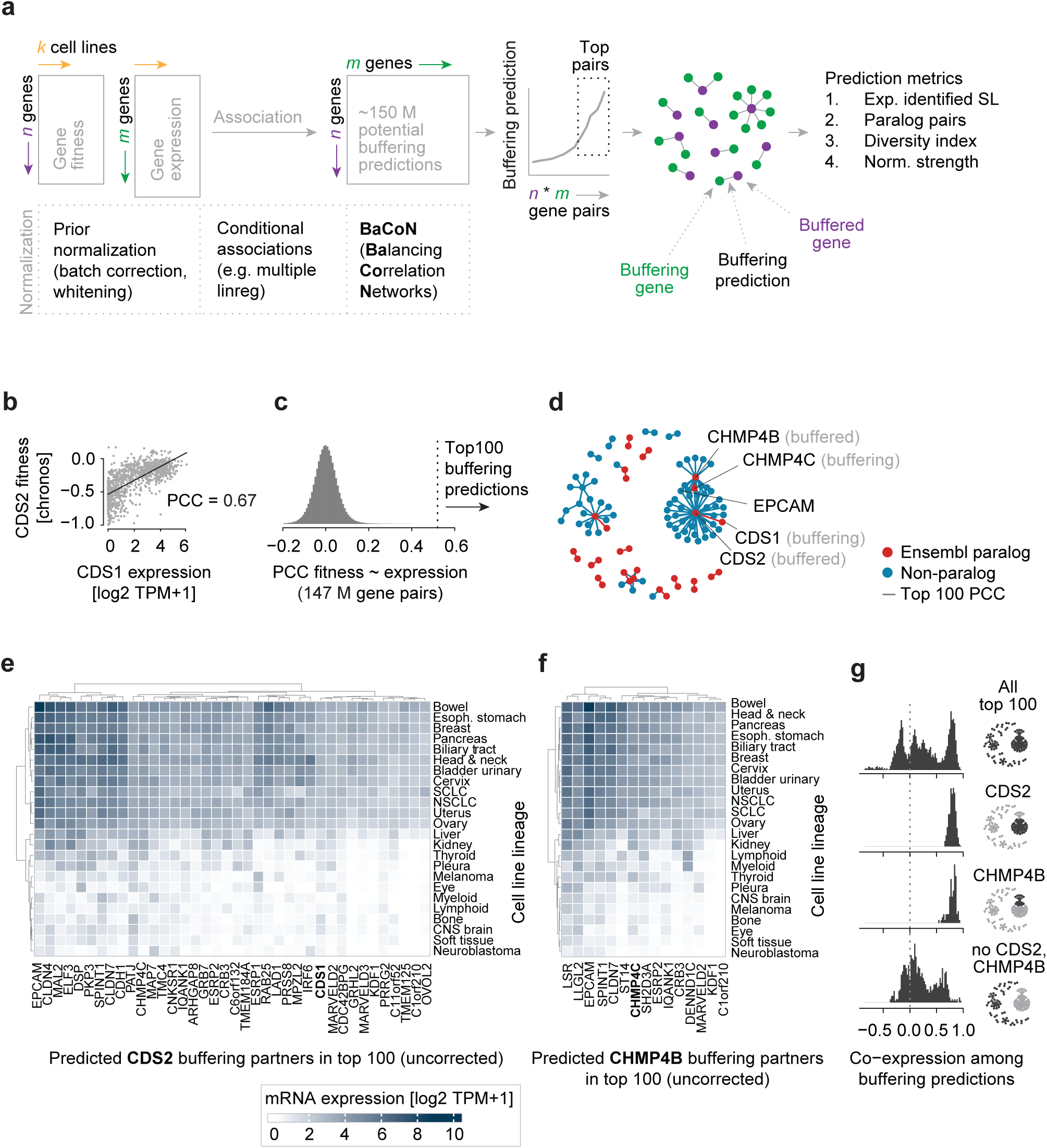
Normalization strategies for predicting gene buffering from the Cancer Dependency Map. **a.** Schematic illustration of normalization strategies where gene buffering is predicted by associating each n gene fitness effect from CRISPR screens with each m gene mRNA expression in k cell lines. N and m can be different but k needs to be matched in both data sets. **b.** Correlation of CDS1 expression with CDS2 perturbation fitness effects (Chronos scores) in 1019 23Q2 cell lines. PCC: Pearson correlation coefficient. **c.** Distribution of all pairwise ∼147 million PCCs between 12,401 potentially buffering genes (CCLE expression data matrix) and 11,885 potentially buffered genes (DepMap Chronos matrix). **d.** Network of the PCC-based top 100 buffering predictions. **e.** and **f.** Mean expression in cell lines derived from the indicated lineages for all genes connected to CDS2 (e.) or CHMP4B (f.) among the top 100 buffering predictions shown in d. **g.** Co-expression of the buffering genes of the top 100 predictions with and without the genes connected to CDS2 and/or CHMP4B. Co-expression is measured as the PCC of log2 TPM+1 values in 1019 cell lines. The network thumbnail (see d) highlights the buffering genes included in a given histogram.

Here, to avoid the risk of unlinking covariation in orthogonal data sets, we developed BaCoN (<u>Ba</u>lanced <u>Co</u>rrelation <u>N</u>etwork), a method to correct correlation-based networks post-hoc (Figure 1a). BaCoN takes a correlation matrix and adjusts the correlation coefficient between each gene pair by balancing it relative to all coefficients each gene partner has with all other genes in the matrix. We assembled four sets of metrics to evaluate systematic buffering prediction performance. Based on those metrics, we benchmark BaCoN and compare it to multiple linear regression and simple correlation coefficients with and without a priori data normalization. We find that BaCoN alone outperforms the other methods including a vastly increased performance over multiple linear regression. We demonstrate that combining the post-hoc method BaCoN with a priori normalization methods further improves performance. Finally, we generate a new, high-quality set of 899 gene pair buffering predictions.

## Results

### Genome-wide prediction of gene buffering

Genome-wide gene buffering can be predicted by defining cell lines where one gene’s function is likely to be absent due to low expression or a loss-of-function mutation and then test if another gene displays more severe fitness effects in just those cell lines (Köferle *et al*, 2022; De Kegel *et al*, 2021; Krieg *et al*, 2024). We systematically evaluate state-of-the-art data normalization methods for predicting gene buffering from the Cancer Dependency Map (DepMap) and present a novel method to further improve such predictions. To build our method, we reasoned that gene buffering predictions, in contrast to co-essentiality or co-expression-derived functional similarity predictions, have two fundamental characteristics. First, genes are likely only buffered by one or a few genes, often paralogs. Second, buffering predictions align two independent data matrices: gene perturbation fitness effects and mRNA expression and independent a priori normalization of those input data potentially unlinks biologically meaningful covariation (Figure 1a, b). Exploiting the few-buffering-gene hypothesis and avoiding the risk of unlinking covariation in independent input data, we developed BaCoN (<u>Ba</u>lanced <u>Co</u>rrelation <u>N</u>etwork), a method to correct correlation-based networks post-hoc (Figure 1a). BaCoN emphasizes specific high pair-wise coefficients by penalizing values for pairs where one or both partners have many similarly high values (see methods for details).

To evaluate gene buffering prediction performance of BaCoN and alternative normalization strategies, we utilized DepMap 23Q2 fitness scores and the associated CCLE expression data (Tsherniak *et al*, 2017; Meyers *et al*, 2017; Barretina *et al*, 2012). DepMap fitness (Chronos) scores measure how much the fitness of each of the 1095 tested cell lines declines upon CRISPR-Cas9-mediated perturbation of each of the roughly 18K protein coding genes. The CCLE expression data contains log2 TPM+1-transformed mRNA expression data for 19193 genes in each of the 1019 cell lines that intersect with the 23Q2 DepMap gene effect data. We restricted our analyses to genes that were expressed at an average log2 TPM+1 > 1 across all cell lines. Overall, this left 12,401 potentially buffering genes (CCLE expression data matrix) and 11,885 potentially buffered genes (DepMap Chronos matrix). Notably, in contrast to co-expression or co-essentiality analysis, (i) the potentially buffered and buffering genes do not have to be identical and (ii) the correlation between gene A and gene B does not equal the correlation between gene B and gene A (Figure 1a). All a priori normalizations, such as batch correction, were performed on these two matrices while the post-hoc method BaCoN was computed on the PCC network derived from those matrices (unless stated otherwise).

### Metrics for evaluation of buffering prediction from the Cancer Dependency Map

To evaluate the performance of BaCoN and various state-of-the-art methods for their ability to predict gene buffering, we defined four metrics. **Firstly**, since systematic, genome-wide searches for buffering gene pairs are expected to be enriched for paralog pairs (De Kegel *et al*, 2021; Köferle *et al*, 2022), we defined the number of Ensembl paralog gene pairs with at least 20% sequence identity as well as the number of Ohnologs among the top buffering predictions as the first metric (Figure S1a) (Singh & Isambert, 2019; Yates *et al*, 2019). **Secondly**, we counted the number of predicted, predicted and validated (Figure S1b) as well as experimentally identified (Figure S1c) synthetic lethal paralog pairs from previous studies (De Kegel *et al*, 2021; Ito *et al*, 2021; Esmaeili Anvar *et al*, 2024; Thompson *et al*, 2021; Dede *et al*, 2020; Gonatopoulos-Pournatzis *et al*, 2020; Parrish *et al*, 2021). Based on those first two sets of metrics, we defined the top 100 and top 1000 gene pairs as the strict and standard buffering predictions (Figure S1d, e) (see methods for details). **Thirdly**, as mentioned above, we reasoned that genes are buffered by one or few other genes, and that the prediction of a large number of buffering genes may be due to confounding factors. In support of this hypothesis, we found that uncorrected correlation of expression and fitness scores predicted some genes to be buffered by a large number of genes among the top 100 buffering predictions of approximately 147 million tested pairs (Figure 1c, d). The fact that (i) among those buffering genes was a paralog partner of the buffered gene, (ii) this paralog partner showed strong tissue-specific expression and (iii) the buffering genes were strongly co-expressed suggested non-buffering false positive prediction of buffering genes when considering correlation without correction. For instance, cell line dependency on *CDS2* showed strong correlation with *CDS1* expression. However, *CDS1* was co-expressed with the epithelial marker gene *EPCAM* and several other genes with epithelial expression patterns (Figure 1e, g). The same was true for *CHMP4B*, which showed a high dependency on the expression of *CHMP4C*, another gene with higher expression in cell lines of epithelial origin (Figure 1f, g). We concluded that those scenarios show high correlation between the gene effect of one gene with the expression of many without a direct buffering relation. To quantify such a bias, we reported the minimal number of genes accounting for 50% of predictions as diversity index between 0 and 1, where 1 indicates that no gene occurs among the predicted pairs more than once (very diverse) (Figure S2a-c). **Lastly**, we assessed how strong a normalization method transformed the original correlation data, with the goal of changing the data as little as possible while removing biases. Specifically, we tested the proportions of pairs within the top buffering predictions after normalization that lacked strong correlation in the original data. Since, before normalization, 788,051 gene pairs showed a z-score-transformed correlation larger than 3 (density of 0.005%), we defined this pool of gene pairs as the acceptable set for the top 100 predictions after normalization. Gene pairs with a z-score-transformed correlation smaller 0, 1, 2 or 3 were all defined as questionable without in-depth exploration (Figure S3a).

### BaCoN outperforms existing methods to predict gene buffering

BaCoN is a post-hoc normalization method for predicting gene buffering by comparing perturbation effects of a gene and mRNA expression of another gene. In its standard implementation, a Pearson correlation coefficient (PCC) matrix between gene effects and mRNA expression is computed first. Overall, applying BaCoN on this PCC matrix (network) improved all four metrics. Among the top 100 buffering predictions BaCoN increased the number of Ensembl and Ohnolog paralog pairs from 16 to 71 and 10 to 53, respectively (Figure 2a). Predicted synthetic lethal (SL) paralog pairs by De Kegel and colleagues increased from 8 to 36 and experimentally identified SL paralog pairs from 7 to 34, 3 to 21, 0 to 5 and 3 to 10, respectively (Figure 2a). BaCoN substantially reduced the bias as shown by an increased diversity index and reduced co-expression of buffering genes (Figure 2b, c). For instance, while *CDS2* and *CHMP4B* were represented in 38 and 15 of the top 100 PCC-based predictions, respectively, the only top 100 pair for *CDS2* was with its paralog partner *CDS1* and *CHMP4B* dropped out of the top 100, with the *CHMP4C* being the strongest partner after BaCoN normalization. At the same time, BaCoN only moderately transformed the PCCs only elevating gene pairs into the top 100 predictions that had a PCC z-score larger 3 (Figure 2d, Figure S3b). Together, this showed that balancing the PCC-based network substantially improved the performance to predict gene buffering. To compare BaCoN performance at removing spurious, likely cell line lineage-driven buffering predictions as described for *CDS2* and *CHPM4B* (Figure 1d-g), we tested several established supervised correction methods. We performed batch correction using ComBat with cell line lineage as batch of the fitness effect and expression matrices separately followed by computing a PCC matrix on the corrected matrices. Counterintuitively, this increased the number of pairs including *CDS2* or *CHMP4B* to 39 and 31, respectively (Figure 2e). In contrast, as expected multiple linear regression between gene effect and mRNA expression where lineage is the covariate reduced the number of pairs including *CDS2* or *CHMP4B* to 15 and 1, respectively (Figure 2e). In conclusion, BaCoN reduced the bias of *CDS2* and *CHMP4B*, likely caused by lineage-driven co-expression of their paralog partner *CDS1* and *CHMP4C*, more effectively than both established methods that correct for lineage in a supervised fashion.

**Figure 2.**
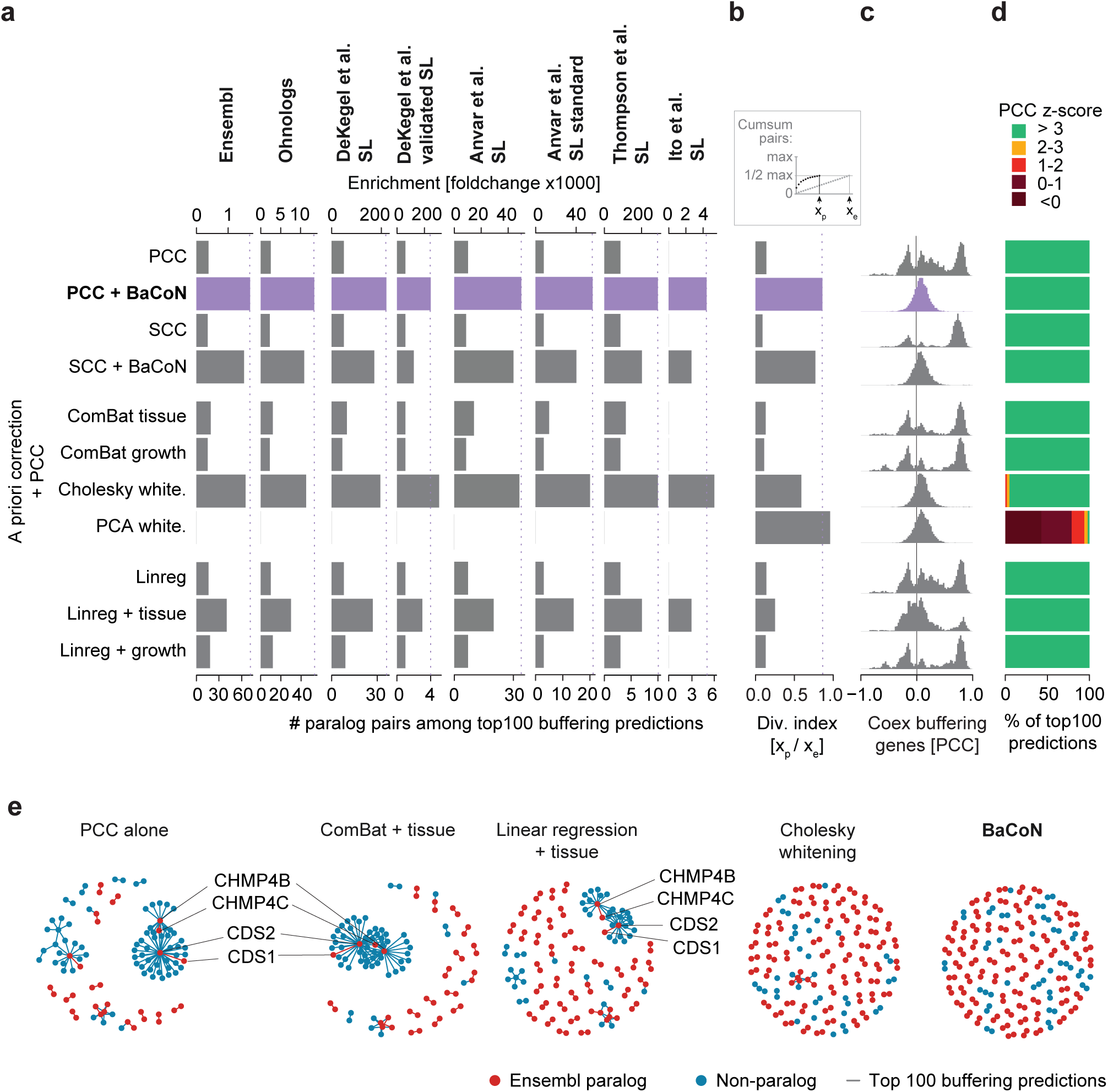
BaCoN outperforms existing methods for predicting gene buffering. **a.** Performance of BaCoN and alternative normalization methods to systematically predict functional buffering. The performance is evaluated using the enrichment (foldchange; fc) and absolute number of sequence-based paralogs, predicted synthetic lethalities (SL) and experimentally identified SLs between paralogs among the top 100 buffering gene pair predictions of each method. Approximately 147 million gene pairs are tested across 1019 different cell lines from the DepMap. PCC: Pearson’s correlation coefficient; SCC: Spearman’s correlation coefficient; Linreg: multiple linear regression with indicated experimental factor as covariate. **b.** Diversity index among the top 100 buffering predictions. **c.** Co-expression among all buffering genes included in the top 100 buffering predictions. Co-expression is measured across the 1019 cell lines used for the buffering predictions. **d.** Strength of data transformation of each method indicated by the numbers of pairs with low z-score-transformed PCCs. This PCC describes the gene fitness effect-vs-expression association and is identical to the PCC shown in panel a. **e.** PCC-based network of the top 100 buffering predictions of five selected methods.

Next, we expanded the comparison of BaCoN to other methods including classical a priori supervised batch correction and noise whitening methods as well as multiple linear regression. We expanded the above-described supervised methods to correct for growth type, a feature distinguishing adherent, suspension or mixed growth of each cell line. However, improvements regarding all metrics were less pronounced than for the lineage-based corrections (Figure 2a-d). Finally, we turned to a method that we and others had independently confirmed to perform very well on co-essentiality networks - data whitening with subsequent computation of pairwise PCCs (Hassan *et al*, 2023; Gheorghe & Hart, 2022; Rahman *et al*, 2021). Data whitening can be done with different sphering methods. We were surprised to see that Cholesky whitening of the gene effect and mRNA expression matrices prior to computing the association indices performed almost as good as BaCoN regarding all metrics (Figure 2a-e). However, Cholesky whitening strongly transforms the data, which leads to several top predictions with low z-score-transformed PCCs, an issue that became dominant when considering larger numbers of buffering predictions (Figure 2d, Figure S4a-l).

Together, BaCoN outperformed supervised correction methods by providing a favorable combination of high paralog pair and known SL predictive performance while only moderately transforming the data.

### Adding BaCoN improves the performance of priori normalization methods

BaCoN is a post-hoc correction method that accounts for biases after association indices such as PCC-based gene buffering predictions have been computed. Since the state-of-the-art alternative methods perform data correction prior to or during association index computation (Figure 1a), we tested if BaCoN could further improve the performance of already normalized scores. Adding BaCoN improved the performance of all methods and across all metrics (Figure 3a-c). The exception is multiple linear regression, the most routinely deployed method for associating gene effect with expression data, since p-values are used instead of association indices (De Kegel *et al*, 2021; Pacini *et al*, 2024). While BaCoN alone already outperformed multiple linear regression (Figure 2a-e), we did not fine-tune BaCoN at this point to consistently improve multiple linear regression association scores (p-values). A priori normalization methods such as batch correction or noise whitening showed increased performance even outperforming PCC plus BaCoN (Figure 3a-c). Especially, normalizing gene effect and expression data using ComBat with defining batches by cell line lineage consistently predicted more buffering paralog pairs and experimentally identified SL paralog genes pairs than BaCoN alone (Figure 3a). In contrast to Cholesky whitening, ComBat batch correction only made predictions for pairs with an original PCC z-score greater 3 (Figure 3c). This suggests that a moderate batch correction prior to BaCoN normalization provides well-adjusted gene buffering predictions without a risk to mistakenly identify gene pairs lacking clear covariance in the original gene effect and expression data. We initially hypothesized that independent normalization of the input data prior to computing association indices could unlink biologically relevant information between the data sets. While PCA-based whitening likely completely unlinked gene fitness effect and expression data, Cholesky whitening showed a surprisingly high number of overall and SL paralog pairs, and Cholesky whitening performance further increased when adding BaCoN (Figure 3a). Notably, the more relaxed list of top 1000 predicted buffering pairs (Figure S1d, e) showed even stronger performance of Cholesky whitening with BaCoN outperforming every other method in every paralog pair and diversity index metric (Figure 3d, e). However, in this list, several pairs were even slightly negatively correlated prior to normalization (Figure 3f). Moreover, the absolute number of pairs with an original PCC z-score smaller than 2 exceeded the number of paralogs and the number of non-paralogs above and below an original PCC z-score of 3 were approximately equal (Figure S4f). This suggested that Cholesky whitening with BaCoN can help predict potential buffering pairs with high sensitivity but partial whitening-based unlinking of gene effect and expression data may also predict potential false buffering pairs.

**Figure 3.**
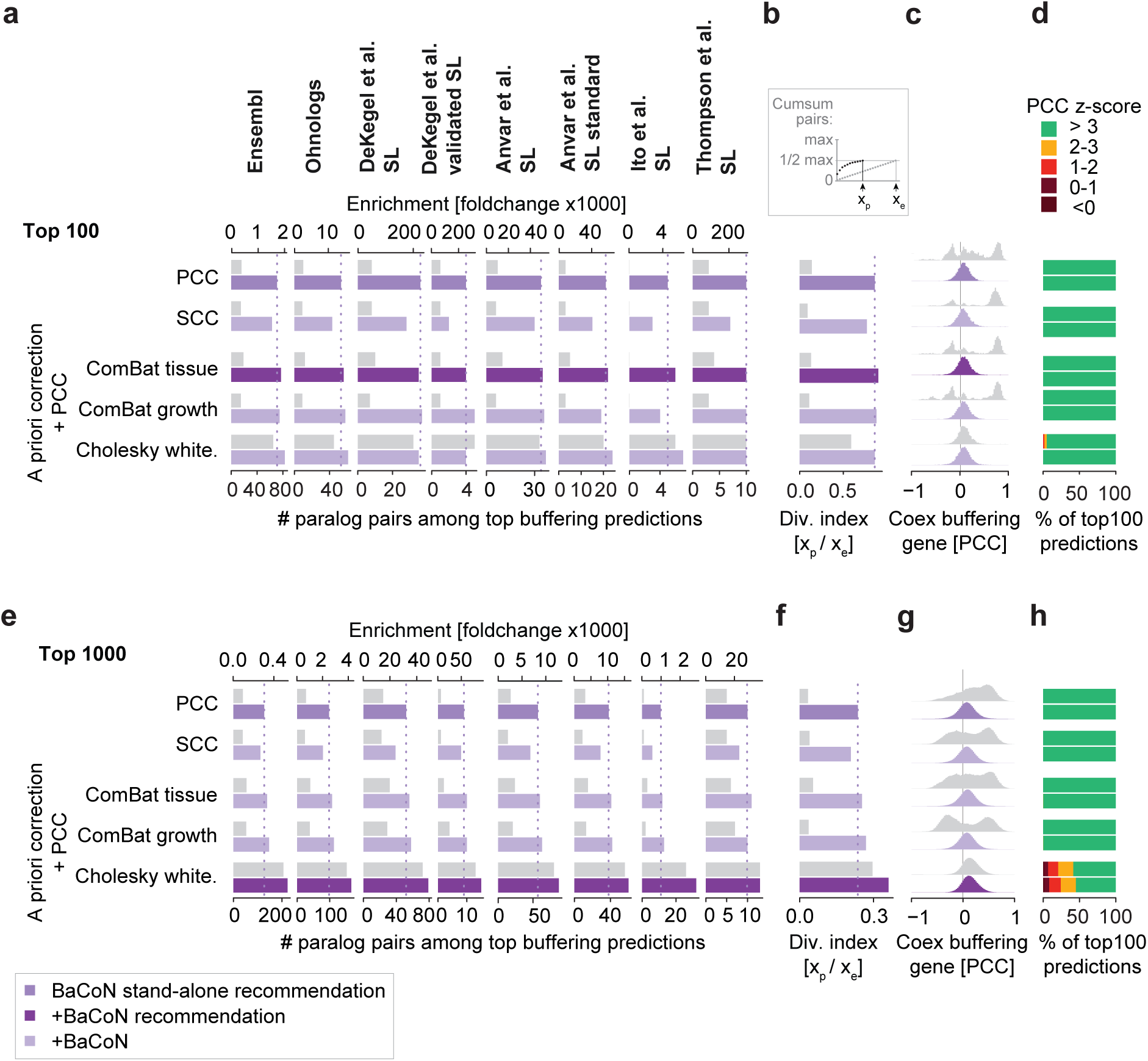
Adding BaCoN improves prediction of gene buffering. **a.** Performance of BaCoN in combination with alternative normalization methods to systematically predict functional buffering. The performance is evaluated using the enrichment (foldchange; fc) and absolute number of sequence-based paralogs, predicted synthetic lethalities (SL) and experimentally identified SLs between paralogs among the top 100 buffering gene pair predictions of each method. Approximately 147 million gene pairs are tested across 1019 different cell lines from the DepMap. PCC: Pearson’s correlation coefficient; SCC: Spearman’s correlation coefficient; Linreg: multiple linear regression with indicated experimental factor as covariate. **b.** Diversity index among the top 100 buffering predictions. **c.** Co-expression among all buffering genes included in the top 100 buffering predictions. Co-expression is measured across the 1019 cell lines used for the buffering predictions**. d.** Strength of data transformation of each method indicated by the numbers of pairs with low z-score-transformed PCCs. This PCC describes the gene fitness effect-vs-expression association and is identical to the PCC shown in panel a. **e.** Performance of BaCoN in combination with alternative normalization methods to systematically predict functional buffering evaluated in a relaxed set of the top 1000 predicted gene pairs. **f.** Diversity index among the top 1000 buffering predictions. **g.** Co-expression among all buffering genes included in the top 1000 buffering predictions. **h.** Strength of data transformation of each method indicated by the numbers of pairs with low z-score-transformed PCCs.

Together, adding BaCoN to established a priori normalization methods can further improve gene buffering predictions. We propose to combine BaCoN with ComBat batch correction for low risk, slightly improved predictive power and BaCoN with Cholesky whitening for higher risk, highly powered predictions of gene buffering.

### BaCoN gains power with growing data sets

The Cancer Dependency Map (DepMap) that we here used for predicting gene buffering has been an ongoing effort mapping gene dependency across various cancer models. More than 1000 different cell lines (models) have been screened and additional screens are performed and released in regular intervals (Boehm *et al*, 2021). To anticipate the potential impact of BaCoN on the future DepMap, we compared its performance with different numbers of screens. We sub-sampled 100, 200, 500 screens each 10 times from the 23Q2 DepMap set of 1019 screens and tested the performance of the PCC, PCC plus BaCoN, multiple linear regression with lineage as covariate, ComBat and ComBat plus BaCoN. When using an already substantial set of 100 cell lines, neither method outperformed the simple PCC (Figure 4a-c; Figure S5a, b). At 200 cell lines, all more sophisticated methods started to outperform the simple PCC, with BaCoN and ComBat plus BaCoN outperforming the other methods and this trend further continued with 500 cell lines (Figure 4a-c; Figure S5c-f). However, only BaCoN was able to increase the number of predicted paralog pairs between 500 cell lines and the full data (Figure 4a-c). Since performance gains did not increase linearly with the number of screens, predictive power to find gene buffering may have started to converge at 1019 screens to a number we might see when testing a future data set.

**Figure 4.**
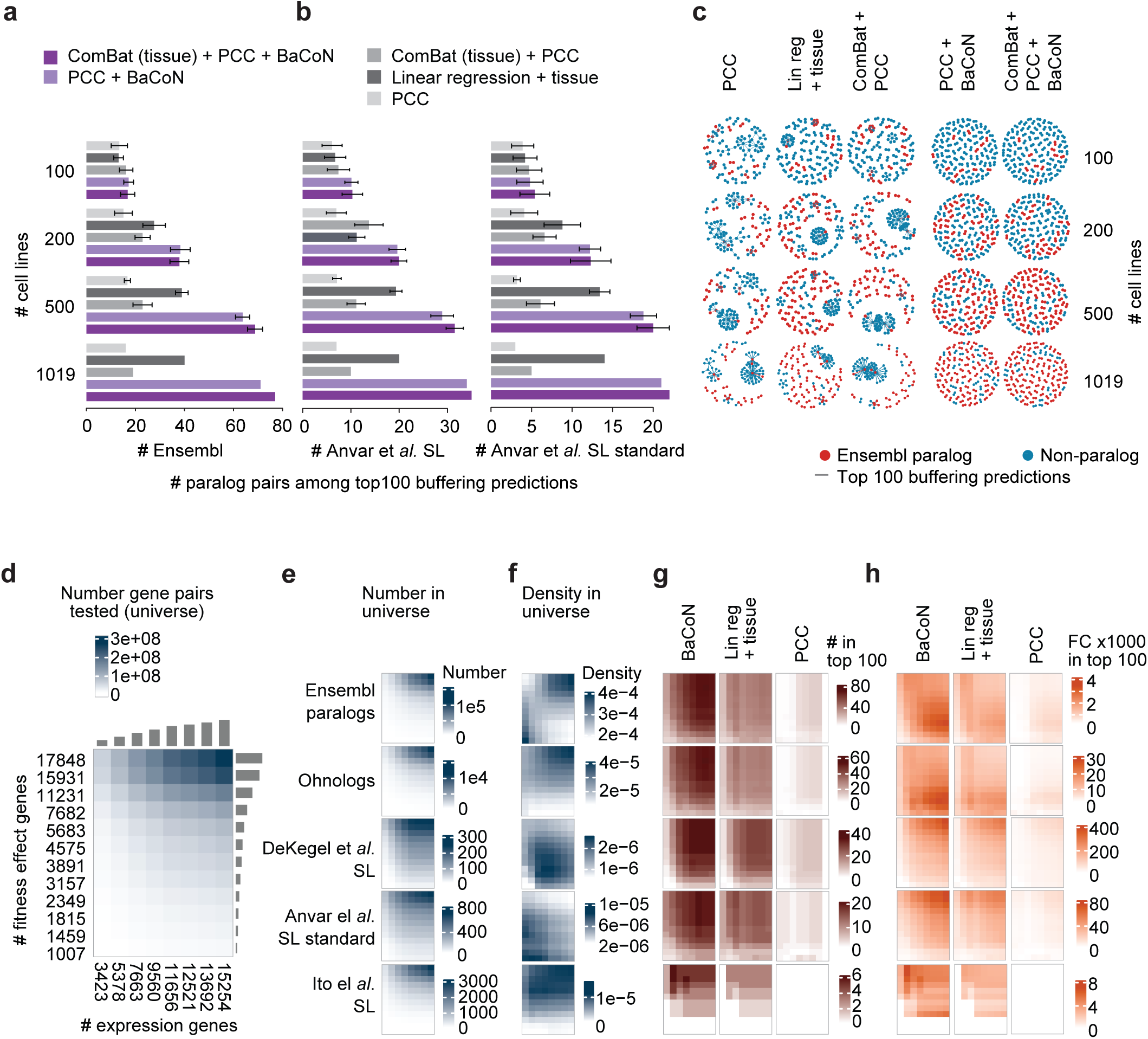
BaCoN improves buffering predictions with a growing number of cell lines and genes tested. **a.** and **b.** Performance of BaCoN and selected alternative normalization methods to predict functional buffering. Performance is quantified as the absolute number of Ensembl paralogs (a.) and experimentally identified SL paralogs (b.) among the top 100 buffering gene pair predictions of each method. Approximately 147 million gene pairs are tested across 100, 200, 500 or 1019 different cell lines from the DepMap. 100, 200 and 500 cell lines were sampled from the 1019 cell lines 10 times and error batch represent the standard deviation of the counts. **c.** PCC-based network of the top 100 buffering predictions of five selected methods. **d.** Gene pair search space when accepting different numbers of genes based on fitness effect and expression thresholds. **e.** Numbers of paralog pairs in the different search spaces. **f.** Density of paralog pairs in the different search spaces**. g.** Numbers of different sets of paralog pairs among the top 100 buffering predictions via BaCoN, multiple linear regression or PCC in the different search spaces. **h.** Enrichment (foldchange; fc) of different sets of paralog pairs among the top 100 buffering predictions via BaCoN, multiple linear regression or PCC in the different search spaces.

We reasoned that future screens added to the DepMap will likely provide cases of a buffering gene losing expression while being widely expressed in the current set of cell lines or the buffered gene displaying currently unseen strong fitness dependency. We tested the impact of such scenarios by testing different numbers of potential buffering and buffered genes by varying the minimal required number of cell lines with fitness effects and loss-of-expression. This spanned input data from 3.4 to 272.3 million gene pairs including 1350 to 107,009 Ensembl paralog and 44 to 11,015 Ohnolog pairs (Figure 4d, e). Among the tested gene pairs, Ensembl paralog or Ohnolog pairs showed higher density when more gene pairs were considered whereas predicted and experimentally identified paralog SLs showed lower density in that area, underlining the investigation bias in current studies (Figure 4f). In those search spaces, similar to multiple linear regression and simple PCC, BaCoN correctly predicted more absolute and relative (foldchange) SL pairs with increasing numbers of potentially buffering genes (Figure 4g, h). When adding more potentially buffered genes, BaCoN tended to benefit from this increased search space more than the other methods. Together, BaCoN particularly gains predictive power in larger data sets, making it a useful tool for future studies in the growing DepMap.

### Characteristics of buffering gene pairs in the human genome

We present BaCoN as a complementary normalization method for predicting gene buffering from the DepMap, and suggest combining BaCoN with Cholesky whitening for sensitive predictions. Deploying this combination of methods, we assembled a high-quality list of 899 buffering gene pairs (see methods for details). We first assessed how those gene buffering predictions are covered by orthogonal data sets and analyses. In contrast to simple standard analysis methods of the same DepMap data, the 899 gene pairs included predictions of *YAP1* buffering its paralog *WWTR1* and *WWTR1* buffering *YAP1*, or *ARID1A* buffering its paralog *ARID1B* and vice versa, none of which showed correlation between gene fitness effect and expression without correction or using multiple linear regression (Figure 5a, b, Figure S3c, f, h). Comparing the predictions with transcriptome-wide Perturb-Seq data (Replogle *et al*, 2022), we found that the expression of the buffering partner was 2.6-fold (*p* = 0.0008) more often up-regulated in response to the perturbation of the buffered partner (Figure 5c). While this mechanistically supports an active buffering mechanism for some predictions, this covered only 15 of the 899 predictions. Since buffering genes must fulfill similar functions to achieve buffering, we tested how the standard approach to predict functional similarity, namely DepMap co-essentiality estimates, cover the 899 buffering predictions. Again, while we observed a highly significant enrichment among our predictions for strong co-essentiality, only 22 pairs were covered (Figure 5d). Finally, similar to previous studies (Esmaeili Anvar *et al*, 2024), paralog pairs with higher sequence similarity were more likely to buffer each other (Figure 5e). However, only around 3% of EMSEMBL pairs with high (>70%) sequence similarity were predicted as buffering (Figure 5f). Together, this highlights the relation between our high-quality list of gene buffering predictions and alternative data sets and methods and stresses the large number of unique observations.

**Figure 5.**
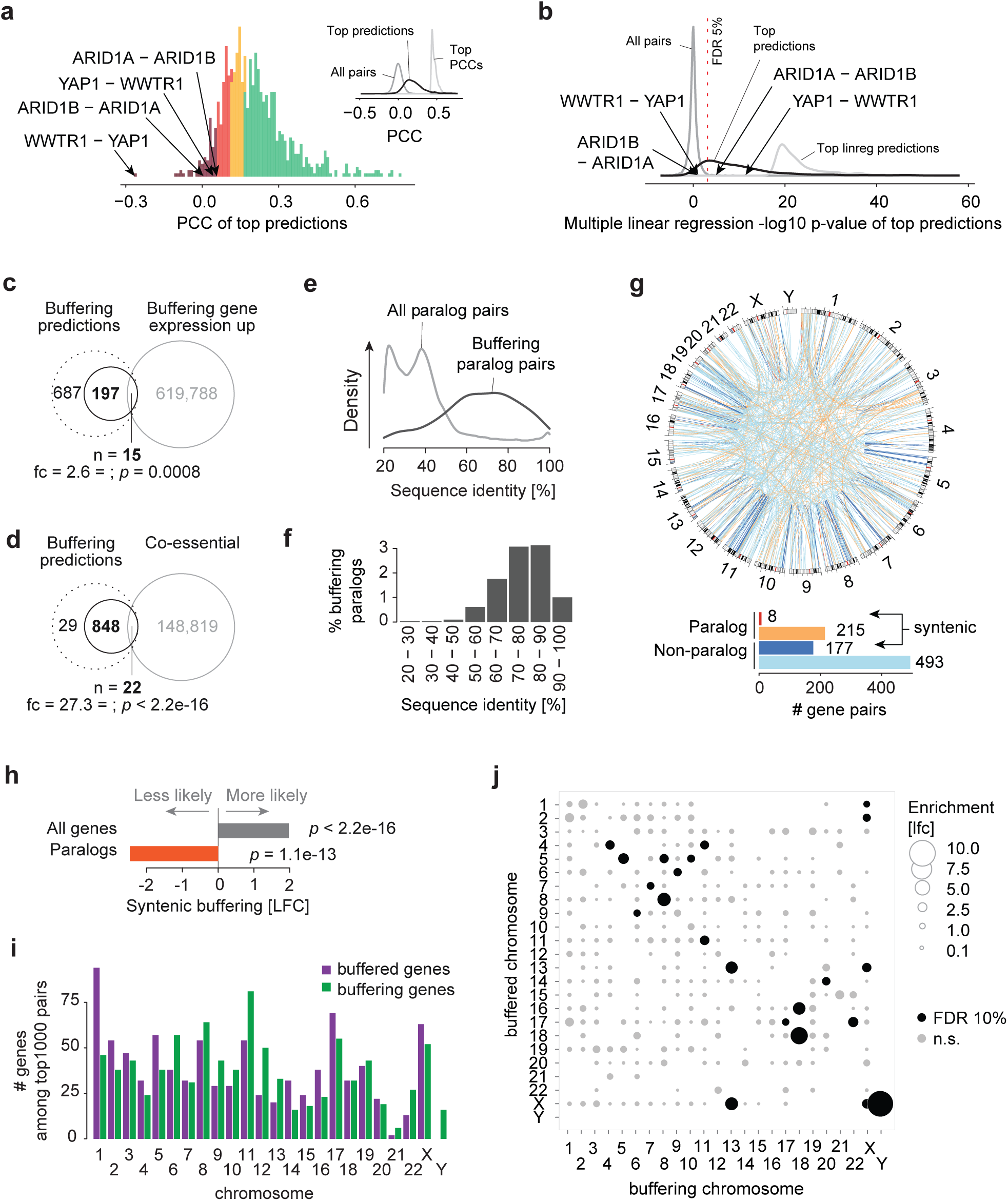
Characteristics of buffering gene pairs in the human genome. 899 high-quality buffering gene pairs were predicted by combining Cholesky whitening followed by BaCoN. **a.** Histogram of fitness effect-vs-expression PCCs of the 899 buffering gene pairs. Bars are colored by the PCC z-score bins (>0, 0-1, 1-2, 2-3, >3). Two paralog pairs were both the AB and BA pair displayed low PCCs and were predicted among the 899 pairs are highlighted. The inset contrasts all 147 million PCCs (grey) with the top 1000 PCCs (light grey) and the PCCs of the 899 high-quality buffering pairs (black). **b.** Multiple linear regression-based buffering prediction scores (-log10 p-val) of all 147 million gene pairs (grey), the top 1000 pairs (light grey) and the 899 high-quality buffering pairs (black). The BH-adjusted p-value (FDR) of 5% is marked. **c.** Overlap of the 899 buffering genes with genes up-regulated when the buffering partner is perturbed in K562 cells (Replogle et al, 2022). For each CRISPRi perturbation in the transcriptome-wide data set by Replogle and colleagues, gene expression signatures were z-score-transformed and to be sensitive, z > 2 was considered up-regulated. The significance of the overlap was tested using a hypergeometric test. **d.** Overlap of the 899 buffering gene pairs with genes showing high co-essentiality in DepMap. Z-score-transformed PCCs larger 4 were considered to show functional similarity. The significance of the overlap was tested using a hypergeometric test. **e.** Sequence identity of Ensembl paralog pairs among the 899 high-quality buffering predictions (n = 223) and the pairs that were not predicted to buffer each other. **f.** Density of high-quality buffering predictions of Ensembl paralogs by sequence identity bin. **g.** Genomic location of paralog and non-paralog high-quality buffering predictions. **h.** Enrichment of both partners of a buffering pair on the same chromosome (syntenic buffering) of Ensembl paralog and non-paralog high-quality buffering predictions. Enrichment was tested for significance using a hypergeometric test. **i.** Number of buffered and buffering genes of the 899 predicted pairs on each chromosome. **j.** Within and between-chromosome density of buffering pairs. Dot size corresponds to foldchange of the density of buffered and buffering genes compared to the global density. Significance was tested by hypergeometric testing with multiple hypothesis correction using the method by Benjamini-Hochberg (BH).

Gene buffering can occur preferentially between different regions in the genome with therapeutic implications (Köferle *et al*, 2022). The 899 buffering predictions were located both within (syntenic) and between chromosomes (Figure 5g). However, while buffering partners were generally 3.9 times more likely to be syntenic (*p* < 2.2e-16) buffering paralog gene pairs were 6.1 times less likely to be on the same chromosome (*p* = 1.1e-13; Figure 5h). Overall, chromosomes harbor different numbers of buffering and buffered genes, with chr1 having the most buffered genes and chr11 showing the most buffering genes (Figure 5i). The buffered-vs-buffering gene imbalance was also detectable for specific chromosome pairs. For instance, chrY genes often buffered chrX genes but not vice versa or chrX buffered chr1 or chr2 more often than expected (Figure 5j). Together, the 899 high-quality buffering predictions covered all chromosomes but showed higher local rates for functional gene buffering.

## Discussion

We present BaCoN (<u>Ba</u>lanced <u>Co</u>rrelation <u>N</u>etwork), a method to correct correlation-based networks for systematically predicting gene buffering from the Cancer Dependency Map (DepMap). BaCoN alone outperforms alternative methods and further increases the performance of predicting gene buffering when combined with a priori data normalization methods. By adding BaCoN to Cholesky whitening, we generate and characterize a high-quality set of 899 gene pair buffering predictions.

Supervised correction of confounding factors such as multiple linear regression represents the standard tool to systematically estimate gene buffering from DepMap gene effect and mRNA expression data (De Kegel *et al*, 2021; Pacini *et al*, 2024; Köferle *et al*, 2022). However, we show that cell lineage causes a major bias among the strongest buffering predictions and that taking into account lineage in multiple linear regression does not remove the bias sufficiently. Interestingly, the simple assumption underlying BaCoN that genes should be buffered by one or a few genes and balancing network degree largely removed this bias in an unsupervised fashion. While similar network balancing strategies have been powerful in other types of molecular networks likely through removing biases (Bass *et al*, 2013; Billmann *et al*, 2016), this strategy matches the expected topology of genome-wide buffering networks.

BaCoN changes prediction score distributions by balancing the strongest predictions in a way that genes with many predicted partners are weakened whereas genes with fewer predicted partners are boosted. This specific post-hoc correction of correlation-based networks only acts on gene pairs with a high level of covariation. In contrast, a priori data normalization methods such as data whitening can remove major axes of variation of the data and emphasize covariation between genes pairs not seen in the original data, which can lead to strongly remodeled prediction score distributions even pinpointing gene pairs with negative correlation in the original data. This is what we coined unlinking of the original input gene effect and expression data and suggest to interpret the resulting scores with caution. When combining Cholesky whitening with BaCoN various metrics showed that the scores outperformed alternative methods, allowing us to define a set of 899 high qualitative gene buffering predictions. This list includes clinically relevant predictions of *YAP1* buffering its paralog *WWTR1* and *WWTR1* buffering *YAP1*, or *ARID1A* buffering its paralog *ARID1B* and vice versa (Frost *et al*, 2023; Helming *et al*, 2014). Notably, all of those four gene pairs displayed a PCC between a z-score of -1 and 1 in the original data, emphasizing that our list of high-quality predictions pinpoint important biological buffering mechanisms that cannot be derived from the DepMap otherwise.

The DepMap has consistently added more cell models and intends to expand past the currently approximately 1100 cell lines targeting more than 10,000 cancer models (Boehm *et al*, 2021). Given that BaCoN particularly excels with an increasing number of screens and genes tested, this suggests that BaCoN can substantially help the search for gene buffering going forward. Notably, while the most comprehensively represented cancer entity in the DepMap counts just over 100 screens, the anticipated increasing number of cell models will likely give rise to tissue-specific, BaCoN-supported gene buffering predictions.

Together, we present BaCoN, a new method to balance correlation networks that use orthogonal inputs such as expression and perturbation data to predict functional buffering between genes without prior knowledge of experimental confounding factors. BaCoN can be combined with existing normalization methods and gains predictive power with increasing numbers of DepMap screens.

## Methods

### NCBI Refseq gene database

Information on the genomic location of genes was retrieved from the NCBI Refseq database (Pruitt *et al*, 2009)(*Homo sapiens*, version GCF_000001405.40_GRCh38.p14). The database was filtered for protein-coding genes.

### Gene effect and expression data

From the DepMap 23Q2 version, CCLE gene expression data ([download](https://depmap.org/portal/download/all/?releasename=DepMap+Public+23Q2&filename=OmicsExpressionProteinCodingGenesTPMLogp1.csv)) as well as Chronos gene fitness effect scores ([download](https://depmap.org/portal/download/all/?releasename=DepMap+Public+23Q2&filename=CRISPRGeneEffect.csv)) were imported. Missing Chronos scores were imputed as gene-wise score means. Genes with a mean expression of less than 1 were removed from the CCLE expression dataset, leaving 12,401 genes in the matrix. The Chronos score matrix was restricted to 11,885 genes that made the gene expression threshold applied to the CCLE data. To allow the computation of association indices, where fitness effects and gene expression was aligned in every cell line, the datasets were reduced to the intersecting 1019 cell lines. Unless stated otherwise, these datasets were used to compute buffering predictions. Cell line metadata on cancer models ([download](https://depmap.org/portal/download/all/?releasename=DepMap+Public+23Q2&filename=Model.csv)) as well as assay conditions ([download](https://depmap.org/portal/download/all/?releasename=DepMap+Public+23Q2&filename=ModelCondition.csv)) was retrieved from the same DepMap version. A list of cell line lineages was defined using OncotreePrimaryDisease for the lineages lung, skin as well as peripheral nervous system, OncotreeLineage classification was used for all other cell lines.

### Defining paralog pairs to evaluate gene buffering predictions

Considering the lack of a comprehensive gold standard set of buffering gene pairs, we assembled a collection of gene pair sets, consisting of human protein-coding Ensembl paralogs and Ohnologs, as well as multiple sets of predicted and/or experimentally validated SL paralog pairs.

#### Ensembl paralogs

A set of 191582 protein-coding paralog pairs was obtained from the *Homo sapiens* Ensembl database (Yates *et al*, 2019), using the BioMart R package (version 2.58.0, data request on 15.04.2024) (Durinck *et al*, 2009). The database request was conducted using the filters “with_hsapiens_paralog” = TRUE and transcript biotype = “protein coding” to select protein coding pairs of human paralogs. The resulting list of paralog pairs was filtered for a minimum sequence identity of 20%.

#### Ohnologs

A set of Ohnolog gene pairs was built based on the Curie database ([download link](http://ohnologs.curie.fr/cgi-bin/DownloadBrowse.cgi?crit=%5BC%5D&org=hsapiens&opt=pairs&wgd=2R)) (Singh & Isambert, 2019), using relaxed filter criteria (outgroup q-score < 0.05 and self-comparison q-score < 0.3). After filtering for protein coding genes, the remaining set contained 7328 pairs (Figure S1a).

#### SL predicted by De Kegel and colleagues

We obtained two sets of paralog pairs from DeKegel et al., 2021 (De Kegel *et al*, 2021). 12 experimentally validated SL paralog pairs ([download link](https://www.cell.com/cms/10.1016/j.cels.2021.08.006/attachment/aa048948-5014-471a-89cf-16b83f522014/mmc2.xlsx)) were defined as ‘DeKegel et al. validated SL’ (Figure S1b). 131 SL paralog predictions defined by a negative A2_status coefficient at 10% FDR were defined as ‘DeKegel et al. SL’ ([download link](https://www.cell.com/cms/10.1016/j.cels.2021.08.006/attachment/8c1dd4f0-6921-4680-a482-5f8199bbb77c/mmc6.csv)).

#### SL pairs defined by Anvar and colleagues

We used two sets of SL paralog pairs based on the work by Anvar and colleagues (Esmaeili Anvar *et al*, 2024). The authors conducted a meta-analysis on five studies that conducted multiplex synthetic lethality screens on paralog pairs, using different Cas12-based systems (Dede *et al*, 2020; Gonatopoulos-Pournatzis *et al*, 2020; Thompson *et al*, 2021; Ito *et al*, 2021; Parrish *et al*, 2021). By re-scoring all pairs tested in these studies, the authors had generated a standard set of 388 paralog pairs. We refer to this set of gene pairs as ‘Anvar et al. SL standard’.

The authors also conducted four screens using two CRISPR/Cas12a multiplex libraries (a prototype library targeting about 2000 paralog families in K-562 and A549 cells, as well as the Inzolia library on A375 and MEL-JUSO cells, targeting about 4000 paralog pairs, triples and quads). In accordance with their approach, we computed the interaction strength (Delta Log Fold Change, dLFC) as the difference between the observed double-knockout LFC of a paralog pair and the sum of single-knock out LFCs. Pairs with a strong negative fitness phenotype (dLFC < -1) in at least one of the screens were defined as hits. We discarded paralog families with more than two genes and used the remaining set of 517 SL paralog pairs as standard. We refer to this set of gene pairs as ‘Anvar et al. SL’ (Figure S1c).

#### SL gene pairs by Ito and colleagues

A set of 1829 paralog pairs was generated based on Gemini scores from the work by Ito and colleagues (Ito *et al*, 2021). We used relaxed filter criteria and required a paralog pair only to show a positive Gemini LFC in at least one of the eleven tested cell lines (FDR < 5%) to be classified as SL (Figure S1c).

#### SL gene pairs by Thompson and colleagues

Thompson and colleagues performed combinatorial CRISPR screens co-perturbing paralog pairs in A375, MeWo and RPE-1 cell lines (Thompson *et al*, 2021). The 27 paralog pairs that were identified as SL in more than one cell line were obtained from this study (Figure S1c).

### Diversity Index

We observed that a small number of genes dominated the top PCC buffering predictions, likely due to confounding with tissue-specific co-expression (Figure 1e-g, Figure S2a). This observation led to the conclusion that low diversity of a prediction set (a small number of genes contributing to a large number of gene pairs) can be used as a marker for biased predictions, and that quantifying the diversity of a prediction set is a useful quality characteristic to compare the predictions made using different correction methods. The diversity index reflects the minimal number of genes contributing to 50% of the pairs in a set of predictions (Figure S2b, c). For very diverse prediction sets (consisting of many independently paired genes) the coefficient is 1. It is computed by first computing the maximum possible number of genes forming 50% of the tested pairs (e.g. 100 genes for a set of 100 predicted pairs). After that, the number of times each gene is observed, starting with the most frequent one, are cumulatively added, until the maximum possible number of genes is reached. The resulting value is divided by the maximum possible number.

### Pearson’s and Spearman’s rank correlation coefficients

To quantify the buffering capacity between a gene pair (**G*_A_* - **G*_B_*), the association between the CCLE expression of **G*_A_* and the Chronos scores of **G*_B_* was quantified. The Pearson’s correlation coefficient (PCC) was used as default association index, using the cor() function in R on the Chronos and gene expression matrix, after aligning the cell lines of both objects to the intersecting 1019. The result is a correlation matrix with the dimensions 12401 x 11885, containing coefficients for each tested gene pair. The Spearman’s rank correlation coefficient (SCC) was computed analogously.

### Multiple linear regression-based gene-gene association

The problem of quantifying the association strength of variables while accounting for effects of known covariates can be addressed using multiple linear regression. For each gene pair of the respective expression and Chronos matrices, we trained a linear model, using the cell line covariates lineage as well as growth pattern (suspension, adherent) as independent variables. For each gene pair **G*_A_*-**G*_B_*, the dependency of the Chronos scores of **G*_B_* on **G*_A_* mRNA expression was computed. The following formulas were tested independently: **G*_B_ Chronos ∼ *G*_A_ Expression*, **G*_B_ Chronos ∼ *G*_A_ Expression + Lineage*, as well as **G*_B_ Chronos ∼ *G*_A_ Expression + Growthpattern*. The predictions were ranked using the negative logarithm of the main predictor variable (**G*_A_ Expression*) p-value as association metric. As the predictor p-value scales independently from the slope of the model, we inverted this index for pairs with a negative slope. This allowed to distinguish between negatively and positively associated pairs. The output of gene pair buffering predictions using linear models is a matrix of negative logarithmic p-values, with the highest positive values describing the strongest association signals.

### Batch correction using ComBat

Batch effects are a common source of confounding in large, multi-experimental datasets, complicating direct comparison of samples of different origin and experimental conditions (Johnson *et al*, 2007). A well-established method to adjust batch effects in microarray-data is a Bayesian framework implemented in the ComBat algorithm. Using the ComBat function from the R sva package (version 3.50.0, (Leek *et al*, 2012)), we adjusted for batch effects within the input datasets. This was done independently for the gene expression data as well as the fitness effects. After that, a correlation matrix was generated from the adjusted data and used to quantify the buffering association strength. As batch information, the cell line lineages of origin as well as on adherent or suspension growth pattern during the cell culture were used.

### Data whitening

Data whitening describes data transformations that normalize covariance and variance (Gheorghe & Hart, 2022). Several methods of data whitening are established, and whitening based on Cholesky decomposition (Cholesky whitening) has been successfully demonstrated to remove bias in co-essentiality networks (Wainberg *et al*, 2021). We independently tested two whitening methods, namely, Cholesky whitening, as well as whitening based on principal component analysis (PCA-whitening). Using the whitening R package (version 1.4.0), both the expression as well as the Chronos data were whitened prior to computing the PCC-correlation matrix. To normalize the covariance between cell lines but not genes, we transposed the input objects prior to whitening and then re-transposed the whitened matrices before computing the correlation matrix.

### BaCoN (<u>Ba</u>lanced <u>Co</u>rrelation <u>N</u>etwork)

BaCoN is deployed on a correlation matrix with the dimensions *n* x *m*, with *n* describing the number of tested genes from the expression dataset and *m* defining the number of genes with Chronos scores. BaCoN balances the correlation of each gene pair *AB* of a matrix based on the fraction of more extreme hits of the genes of the respective pair. For each gene pair *AB* of the correlation matrix with *ρ_AB_ ≥ 0*, first a correction factor *k* is subtracted from the correlation coefficient *ρ_AB_*. After that, the density of all adjacent and more extreme correlation scores is calculated and subtracted from 1, resulting in high BaCoN scores for gene pairs with few more extreme partners:

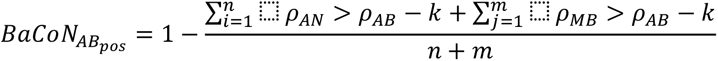

For each pair *AB* with a negative correlation (*ρ_AB_ < 0*), BaCoN scores for negatively correlating gene pairs are generated analogously, with inverted signs. The correction factor *k* is added to *ρ_AB_*, and the density of lower correlations of the pair is added to -1:

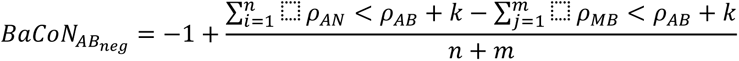

The result is a matrix with the same dimensions as the input matrix.

### Random subsets

Reliable predictions on gene buffering require a sufficient number of cell lines to associate the expression of one gene with the fitness scores of another gene. We quantified the impact of varying the number of cell lines used on predicting buffering to estimate the impact of BaCoN on the future DepMap and to benchmark the buffering prediction capacity of the best-performing methods (PCC, PCC + BaCoN, ComBat (tissue) + PCC, ComBat (tissue) + PCC + BaCoN, multiple linear regression with tissue as covariate). The 1019 overlapping cell lines between the gene expression and Chronos matrices were randomly subsampled. Sets of the sizes 100, 200 as well as 500 were generated, with 10 replicates each. We used the subsampled expression and Chronos matrices to generate association matrices using the previously described methods, keeping the number of tested gene pairs constant.

### Dynamic gene space

Conceptually, we expect that buffering predictions require the buffering partner to be expressed in a sufficient number of cell lines and the knockout of the buffered partner to have a certain impact on cell fitness. To test how removing genes with low expression or essentiality signals influences predictions, we used two types of thresholds that were applied on the gene expression and chronos score data, respectively. The expression thresholds (10, 30, 60, 100, 300, 600, 900, 1000) were used to remove genes that do not show an expression (log2 TPM+1) of *≥* 3 in at least the respective number of cell lines, removing genes with low expression signal. The Chronos thresholds (-0.2, -0.25, -0.3, -0.35, -0.4, -0.45, -0.5, -0.6, -0.8, -1.0, -1.2 and -1.5) were used to select only genes that did show the respective essentiality in a minimum of 30 cell lines. This way, genes with low essentiality across most of the cell lines were removed. Each of the 8 subset gene expression matrices was combined with each of the 12 subset Chronos matrices, resulting in a total of 96 prediction matrices of different dimensions (Figure 4d). The resulting matrices were used to predict buffering, using the overlapping 1019 cell lines. We tested the methods PCC, multiple linear regression with lineage as covariate as well as BaCoN. The number of gene pairs in the respective universe was computed as the product of the number of all genes in the expression data (potential buffering) and all potentially buffered genes (Chronos scores) that were not filtered by applying the filters. Using the previously described reference gene pair sets, the paralog density was computed as the number of gene pair hits divided by the number of total gene pairs of a matrix. The paralog enrichment was computed as the density of paralog hits among the top 100 predictions, divided by the paralog density.

### High-confidence buffering gene pair set

We aimed to generate a larger set of high-quality buffering predictions. Based on our finding that the top 1000 predictions generated by combining Cholesky whitening with BaCoN outperformed the tested competing methods in terms of paralog prediction performance as well as prediction diversity (Figure S4), we based a high-quality buffering prediction set on this set of gene pairs. As described previously, we computed a PCC-based correlation matrix from Cholesky-whitened fitness effect as well as gene expression data. BaCoN was then applied on the resulting PCC matrix. We sorted the pairs decreasingly by their BaCoN score as well as PCC and selected the top 1000 pairs. After filtering self-buffering pairs (pairs with matching buffering and buffered gene), 899 pairs remained in the set.

## Code availability

The BaCoN R package can be obtained from https://github.com/billmannlab/BaCoN/. A Python command line implementation can be obtained from the same repository or https://github.com/billmannlab/pyBaCoN/.

## Acknowledgement

We thank all members from the Billmann lab and the Institute of Human Genetics at the University of Bonn and University Hospital Bonn for helpful discussions.

## Competing interest

The authors declare to competing interest.

---

**Figure S1.**
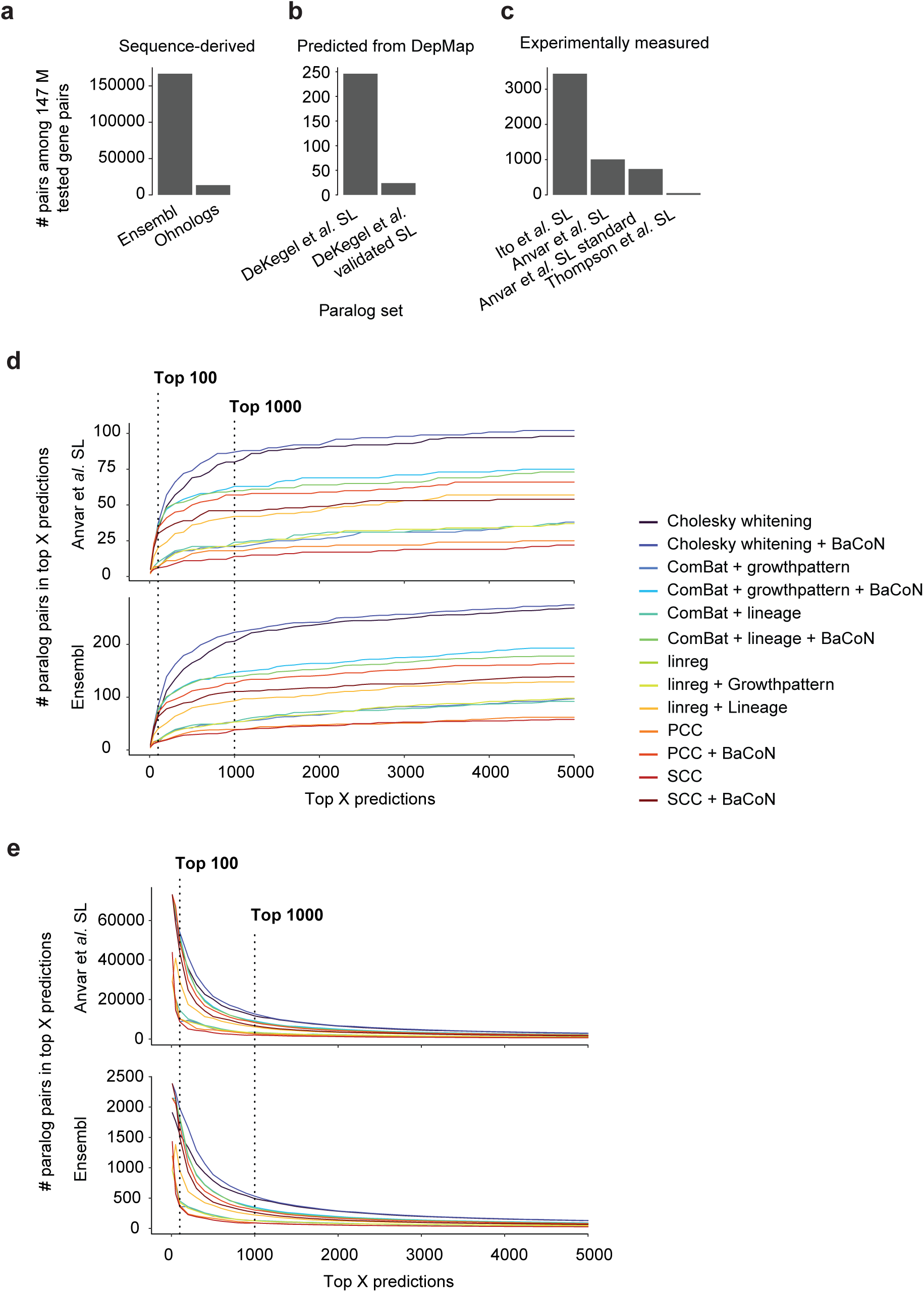
Number of paralog gene pairs in the tested universe of 147 million gene pairs and their numbers among the top 5000 predictions. **a.** Number of Ensembl and Ohnolog paralog pairs in the universe. **b.** Number of predicted synthetic lethal (SL) gene pairs in the universe. **c.** Number of experimentally identified SL paralog pairs in different studies. **d.** Number of Ensembl paralog pairs and experimentally identified SL pairs among the up to 5000 top predictions using different normalization methods alone or in combination. The top 100 and 1000 predicted pairs are marked. **e.** Foldchange (number over all predicted) of Ensembl paralog pairs and experimentally identified SL pairs among the up to 5000 top predictions using different normalization methods alone or in combination. The top 100 and 1000 predicted pairs are marked.

**Figure S2.**
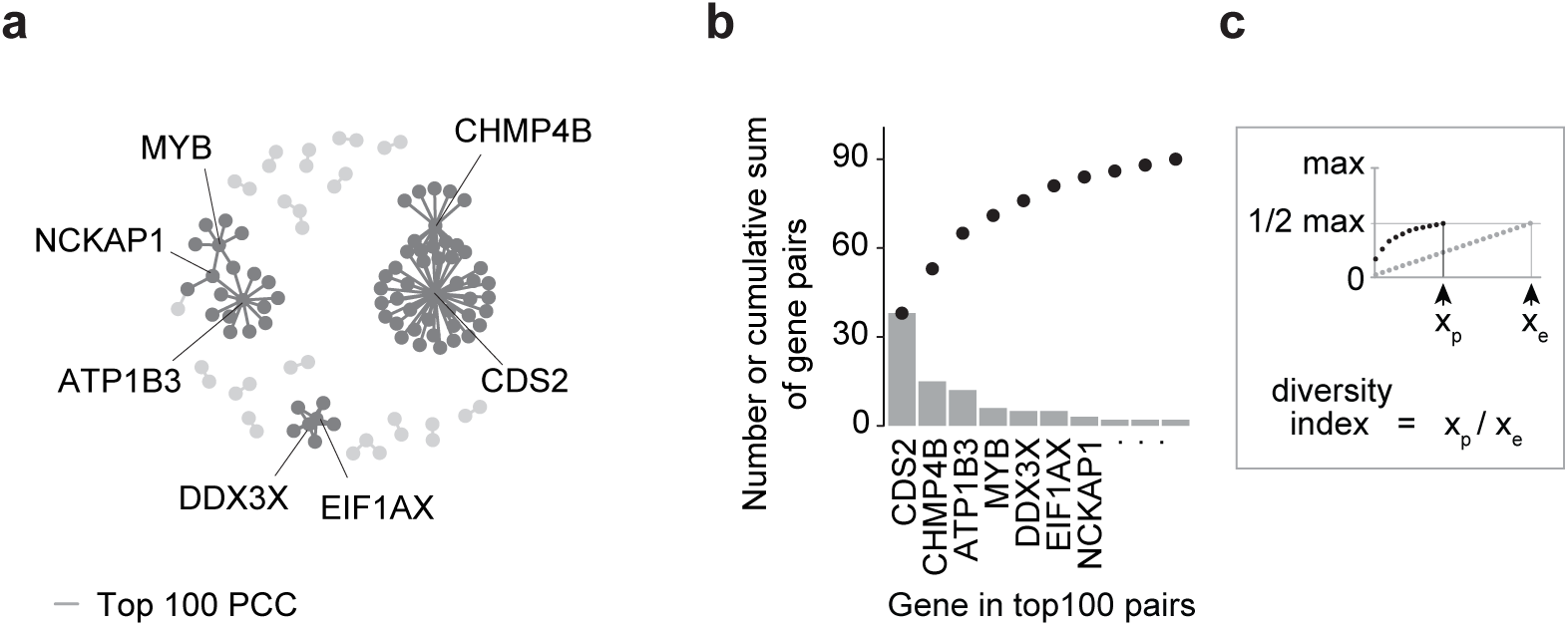
The diversity index among the top buffering predictions. **a.** Network of the PCC-based top 100 buffering predictions. Genes that are predicted to be buffered by at least three genes are labeled. **b.** Number of top 100 pairs a gene is involved in (bars) and cumulative sum of pairs (dots). Genes that are predicted to be buffered by at least three genes are labeled. **c.** Illustration of the diversity index when defining x_p_ (number of genes) at the half maximum number, where x_e_ is the maximum expected number of e.g. 200 genes in top 100 pairs.

**Figure S3.**
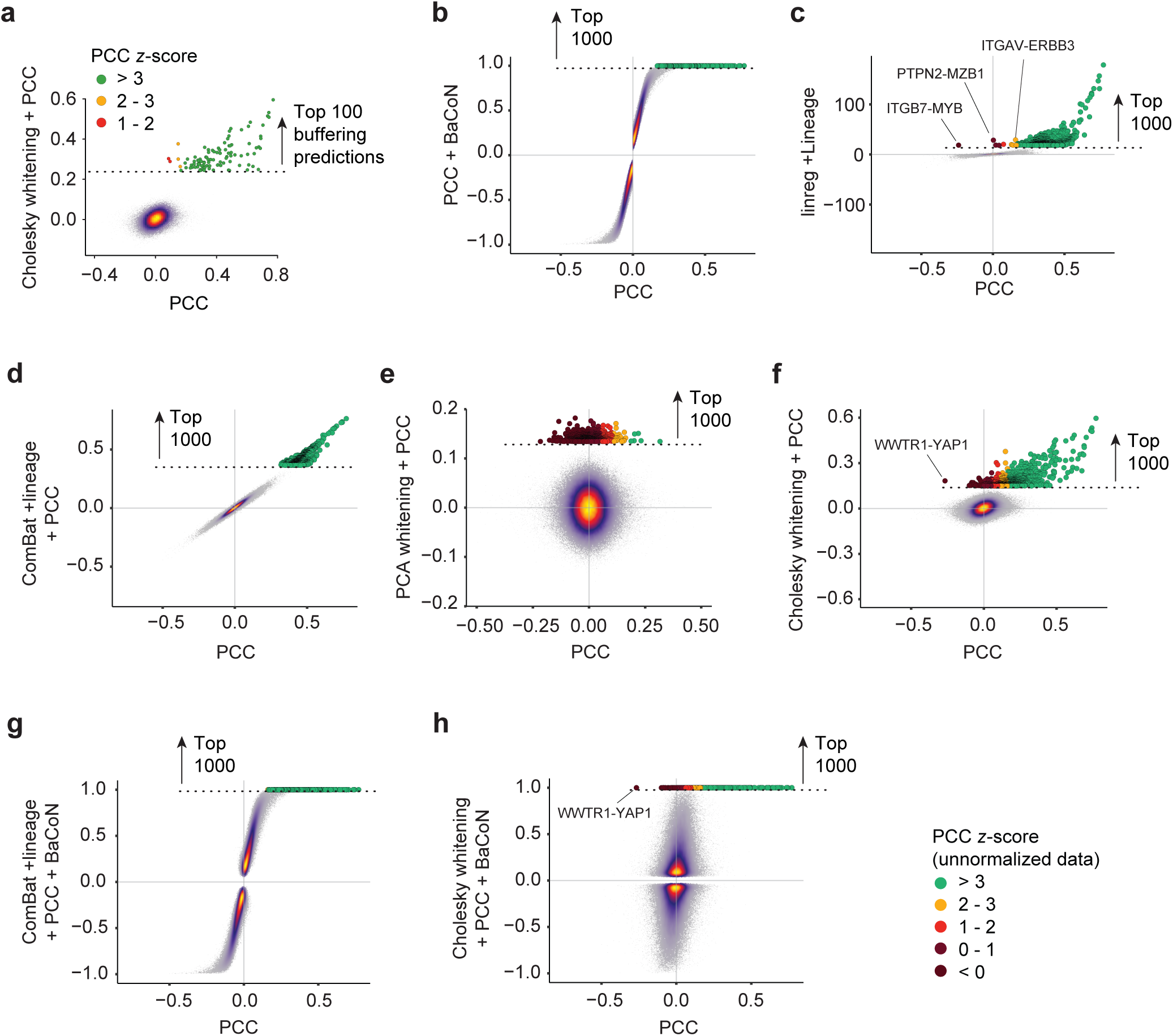
Transformation of buffering prediction scores when utilizing different methods to normalize gene effect and expression data. All methods have been performed on the approximately 147 million gene pairs. **a.** Cholesky whitening plus PCC-based compared to PCC only. All scores are shown as heatscatter. The top 100 Cholesky whitening plus PCC predictions are shown larger and color-coded by z-score bin of the PCC only scores. **b. – h.** Selected normalization method-based predictions compared to PCC only. All scores are shown as heatscatter. The top 1000 corrected predictions are shown larger and color-coded by z-score bin of the PCC only scores. Selected strongly transformed gene pairs are labeled.

**Figure S4.**
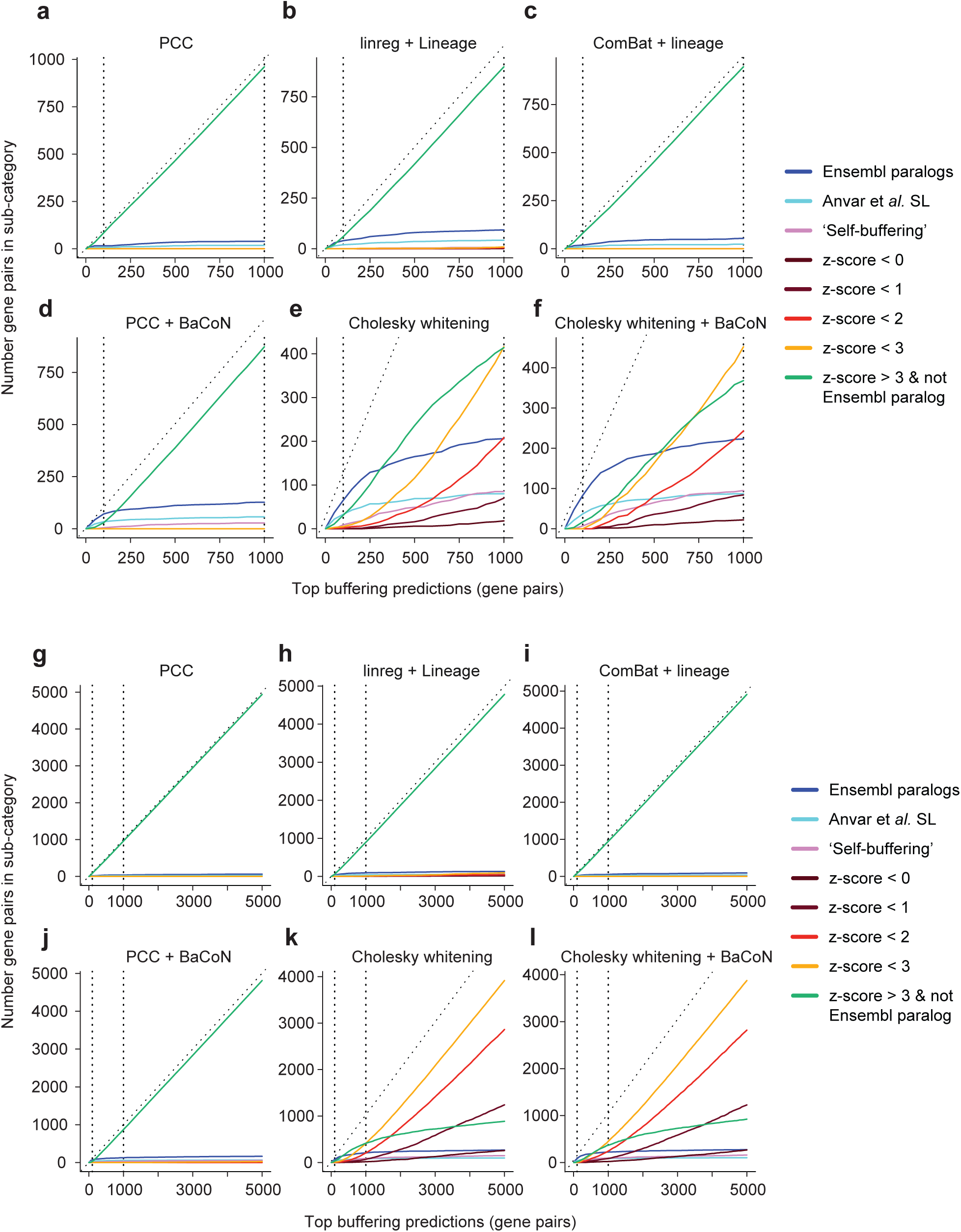
Number of gene pairs in several sub-categories among the top predictions using different normalization methods. Sub-categories are the number of Ensembl gene pairs, experimentally identified SL gene pairs, number of highly correlated pairs that are not Ensembl paralogs, number of ‘self-buffering’ and the number of pairs with low correlation in the original data at different thresholds. **a. – f.** Number of gene pairs of each sub-category among the up to top 1000 predictions. The dotted diagonal is the maximum value a sub-category can reach. The high-confidence (top 100) and standard (top 1000) prediction thresholds are marked. **g. – l.** Number of gene pairs of each sub-category among the up to top 5000 predictions.

**Figure S5.**
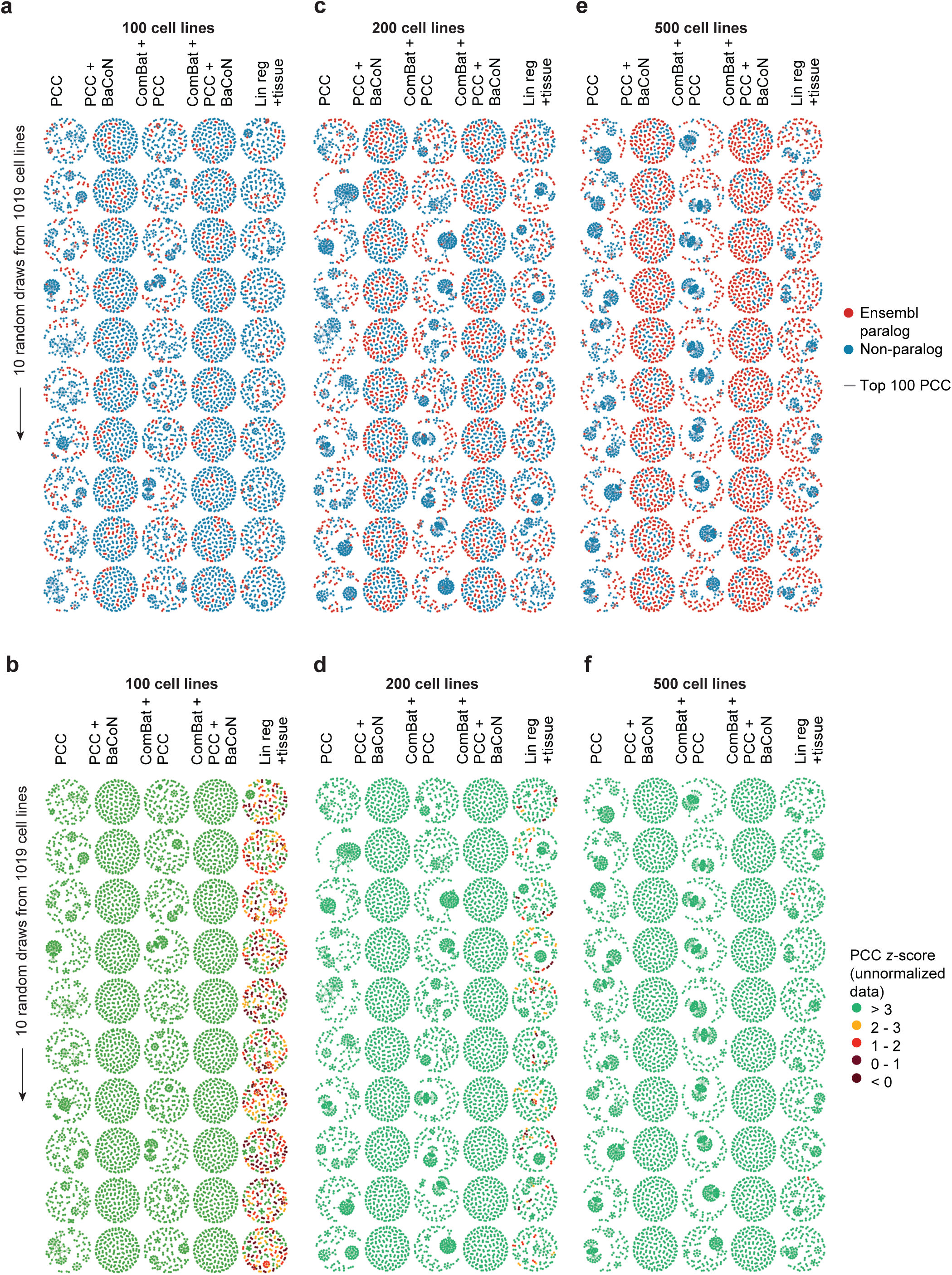
Network visualization of the top 100 buffering predictions using 10 random samples from the 1019 DepMap cell lines. This figure accompanies Figure 4c providing all 10 random samples from which a representative network was chosen. **a.** Top 100 buffering prediction networks from 10 samples of 100 out of 1019 cell lines. Ensembl paralogs and non-paralogs are colored. **b.** Top 100 buffering prediction networks from 10 samples of 100 out of 1019 cell lines. Low correlations in the original uncorrected data are colored. **c.** Top 100 buffering prediction networks from 10 samples of 200 out of 1019 cell lines. Ensembl paralogs and non-paralogs are colored. **d.** Top 100 buffering prediction networks from 10 samples of 200 out of 1019 cell lines. Low correlations in the original uncorrected data are colored. **e.** Top 100 buffering prediction networks from 10 samples of 500 out of 1019 cell lines. Ensembl paralogs and non-paralogs are colored. **f.** Top 100 buffering prediction networks from 10 samples of 500 out of 1019 cell lines. Low correlations in the original uncorrected data are colored.

## Notes

https://github.com/billmannlab/BaCoN/

### Competing Interest Statement

The authors have declared no competing interest.

## References

Aregger M, Lawson KA, Billmann M, Costanzo M, Tong AHY, Chan K, Rahman M, Brown KR, Ross C, Usaj M, et al (2020) Systematic mapping of genetic interactions for de novo fatty acid synthesis identifies C12orf49 as a regulator of lipid metabolism. Nat Metab 2: 499–513

Barretina J, Caponigro G, Stransky N, Venkatesan K, Margolin AA, Kim S, Wilson CJ, Lehár J, Kryukov GV, Sonkin D, et al (2012) The Cancer Cell Line Encyclopedia enables predictive modelling of anticancer drug sensitivity. Nature 483: 603–607

Bass JIF, Diallo A, Nelson J, Soto JM, Myers CL & Walhout AJM (2013) Using networks to measure similarity between genes: association index selection. Nat Methods 10: 1169–1176

Behan FM, Iorio F, Picco G, Gonçalves E, Beaver CM, Migliardi G, Santos R, Rao Y, Sassi F, Pinnelli M, et al (2019) Prioritization of cancer therapeutic targets using CRISPR–Cas9 screens. Nature 568: 511–516

Billmann M, Chaudhary V, ElMaghraby MF, Fischer B & Boutros M (2018) Widespread Rewiring of Genetic Networks upon Cancer Signaling Pathway Activation. Cell Syst 6: 52–64.e4

Billmann M, Horn T, Fischer B, Sandmann T, Huber W & Boutros M (2016) A genetic interaction map of cell cycle regulators. Mol Biol Cell 27: 1397–1407

Boehm JS, Garnett MJ, Adams DJ, Francies HE, Golub TR, Hahn WC, Iorio F, McFarland JM, Parts L & Vazquez F (2021) Cancer research needs a better map. Nature 589: 514–516

Boyle EA, Pritchard JK & Greenleaf WJ (2018) High-resolution mapping of cancer cell networks using co-functional interactions. Mol Syst Biol 14: e8594

Collins SR, Miller KM, Maas NL, Roguev A, Fillingham J, Chu CS, Schuldiner M, Gebbia M, Recht J, Shales M, et al (2007) Functional dissection of protein complexes involved in yeast chromosome biology using a genetic interaction map. Nature 446: 806–810

Costanzo M, Baryshnikova A, Bellay J, Kim Y, Spear ED, Sevier CS, Ding H, Koh JLY, Toufighi K, Mostafavi S, et al (2010) The Genetic Landscape of a Cell. Science 327: 425–431

Costanzo M, VanderSluis B, Koch EN, Baryshnikova A, Pons C, Tan G, Wang W, Usaj M, Hanchard J, Lee SD, et al (2016) A global genetic interaction network maps a wiring diagram of cellular function. Science 353: aaf1420–aaf1420

De Kegel B, Quinn N, Thompson NA, Adams DJ & Ryan CJ (2021) Comprehensive prediction of robust synthetic lethality between paralog pairs in cancer cell lines. Cell Syst 12: 1144–1159.e6

De Kegel B & Ryan CJ (2023) Paralog dispensability shapes homozygous deletion patterns in tumor genomes. Mol Syst Biol 19: e11987

Dede M, McLaughlin M, Kim E & Hart T (2020) Multiplex enCas12a screens detect functional buffering among paralogs otherwise masked in monogenic Cas9 knockout screens. Genome Biol 21: 262

Dempster JM, Boyle I, Vazquez F, Root DE, Boehm JS, Hahn WC, Tsherniak A & McFarland JM (2021) Chronos: a cell population dynamics model of CRISPR experiments that improves inference of gene fitness effects. Genome Biol 22: 343

DeWeirdt PC, Sanson KR, Sangree AK, Hegde M, Hanna RE, Feeley MN, Griffith AL, Teng T, Borys SM, Strand C, et al (2021) Optimization of AsCas12a for combinatorial genetic screens in human cells. Nat Biotechnol 39: 94–104

Durinck S, Spellman PT, Birney E & Huber W (2009) Mapping identifiers for the integration of genomic datasets with the R/Bioconductor package biomaRt. Nat Protoc 4: 1184–1191

Eisen MB, Spellman PT, Brown PO & Botstein D (1998) Cluster analysis and display of genome-wide expression patterns. Proc Natl Acad Sci 95: 14863–14868

Esmaeili Anvar N, Lin C, Ma X, Wilson LL, Steger R, Sangree AK, Colic M, Wang SH, Doench JG & Hart T (2024) Efficient gene knockout and genetic interaction screening using the in4mer CRISPR/Cas12a multiplex knockout platform. Nat Commun 15: 3577

Farmer H, McCabe N, Lord CJ, Tutt ANJ, Johnson DA, Richardson TB, Santarosa M, Dillon KJ, Hickson I, Knights C, et al (2005) Targeting the DNA repair defect in BRCA mutant cells as a therapeutic strategy. Nature 434: 917–921

Fischer B, Sandmann T, Horn T, Billmann M, Chaudhary V, Huber W & Boutros M (2015) A map of directional genetic interactions in a metazoan cell. eLife 4: e05464

Frost TC, Gartin AK, Liu M, Cheng J, Dharaneeswaran H, Keskin DB, Wu CJ, Giobbie-Hurder A, Thakuria M & DeCaprio JA (2023) YAP1 and WWTR1 expression inversely correlates with neuroendocrine markers in Merkel cell carcinoma. J Clin Invest 133: e157171

Gheorghe V & Hart T (2022) Optimal construction of a functional interaction network from pooled library CRISPR fitness screens Systems Biology

Gonatopoulos-Pournatzis T, Aregger M, Brown KR, Farhangmehr S, Braunschweig U, Ward HN, Ha KCH, Weiss A, Billmann M, Durbic T, et al (2020) Genetic interaction mapping and exon-resolution functional genomics with a hybrid Cas9–Cas12a platform. Nat Biotechnol 38: 638–648

Hassan AZ, Ward HN, Rahman M, Billmann M, Lee Y & Myers CL (2023) Dimensionality reduction methods for extracting functional networks from large-scale CRISPR screens. Mol Syst Biol 19: e11657

Helming KC, Wang X, Wilson BG, Vazquez F, Haswell JR, Manchester HE, Kim Y, Kryukov GV, Ghandi M, Aguirre AJ, et al (2014) ARID1B is a specific vulnerability in ARID1A-mutant cancers. Nat Med 20: 251–254

Iorio F, Behan FM, Gonçalves E, Bhosle SG, Chen E, Shepherd R, Beaver C, Ansari R, Pooley R, Wilkinson P, et al (2018) Unsupervised correction of gene-independent cell responses to CRISPR-Cas9 targeting. BMC Genomics 19: 604

Ito T, Young MJ, Li R, Jain S, Wernitznig A, Krill-Burger JM, Lemke CT, Monducci D, Rodriguez DJ, Chang L, et al (2021) Paralog knockout profiling identifies DUSP4 and DUSP6 as a digenic dependence in MAPK pathway-driven cancers. Nat Genet 53: 1664–1672

Johnson WE, Li C & Rabinovic A (2007) Adjusting batch effects in microarray expression data using empirical Bayes methods. Biostatistics 8: 118–127

Köferle A, Schlattl A, Hörmann A, Thatikonda V, Popa A, Spreitzer F, Ravichandran MC, Supper V, Oberndorfer S, Puchner T, et al (2022) Interrogation of cancer gene dependencies reveals paralog interactions of autosome and sex chromosome-encoded genes. Cell Rep 39: 110636

Krieg S, Rohde T, Rausch T, Butthof L, Wendler-Link L, Eckert C, Breuhahn K, Galy B, Korbel J, Billmann M, et al (2024) Mitoferrin2 is a synthetic lethal target for chromosome 8p deleted cancers. Genome Med 16: 83

Kryukov GV, Wilson FH, Ruth JR, Paulk J, Tsherniak A, Marlow SE, Vazquez F, Weir BA, Fitzgerald ME, Tanaka M, et al (2016) *MTAP* deletion confers enhanced dependency on the PRMT5 arginine methyltransferase in cancer cells. Science 351: 1214–1218

Leek JT, Johnson WE, Parker HS, Jaffe AE & Storey JD (2012) The sva package for removing batch effects and other unwanted variation in high-throughput experiments. Bioinformatics 28: 882–883

Lord CJ, Quinn N & Ryan CJ (2020) Integrative analysis of large-scale loss-of-function screens identifies robust cancer-associated genetic interactions. eLife 9: e58925

Meyers RM, Bryan JG, McFarland JM, Weir BA, Sizemore AE, Xu H, Dharia NV, Montgomery PG, Cowley GS, Pantel S, et al (2017) Computational correction of copy number effect improves specificity of CRISPR–Cas9 essentiality screens in cancer cells. Nat Genet 49: 1779–1784

Oser MG, Fonseca R, Chakraborty AA, Brough R, Spektor A, Jennings RB, Flaifel A, Novak JS, Gulati A, Buss E, et al (2019) Cells Lacking the *RB1* Tumor Suppressor Gene Are Hyperdependent on Aurora B Kinase for Survival. Cancer Discov 9: 230–247

Pacini C, Duncan E, Gonçalves E, Gilbert J, Bhosle S, Horswell S, Karakoc E, Lightfoot H, Curry E, Muyas F, et al (2024) A comprehensive clinically informed map of dependencies in cancer cells and framework for target prioritization. Cancer Cell: S1535610823004440

Pan J, Kwon JJ, Talamas JA, Borah AA, Vazquez F, Boehm JS, Tsherniak A, Zitnik M, McFarland JM & Hahn WC (2022) Sparse dictionary learning recovers pleiotropy from human cell fitness screens. Cell Syst 13: 286–303.e10

Parrish PCR, Thomas JD, Gabel AM, Kamlapurkar S, Bradley RK & Berger AH (2021) Discovery of synthetic lethal and tumor suppressor paralog pairs in the human genome. Cell Rep 36: 109597

Pruitt KD, Tatusova T, Klimke W & Maglott DR (2009) NCBI Reference Sequences: current status, policy and new initiatives. Nucleic Acids Res 37: D32–D36

Rahman M, Billmann M, Costanzo M, Aregger M, Tong AHY, Chan K, Ward HN, Brown KR, Andrews BJ, Boone C, et al (2021) A method for benchmarking genetic screens reveals a predominant mitochondrial bias. Mol Syst Biol 17

Replogle JM, Saunders RA, Pogson AN, Hussmann JA, Lenail A, Guna A, Mascibroda L, Wagner EJ, Adelman K, Lithwick-Yanai G, et al (2022) Mapping information-rich genotype-phenotype landscapes with genome-scale Perturb-seq. Cell 185: 2559–2575.e28

Ryan CJ, Devakumar LPS, Pettitt SJ & Lord CJ (2023a) Complex synthetic lethality in cancer. Nat Genet 55: 2039–2048

Ryan CJ, Mehta I, Kebabci N & Adams DJ (2023b) Targeting synthetic lethal paralogs in cancer. Trends Cancer 9: 397–409

Singh PP & Isambert H (2019) OHNOLOGS v2: a comprehensive resource for the genes retained from whole genome duplication in vertebrates. Nucleic Acids Res: gkz909

Thompson NA, Ranzani M, Van Der Weyden L, Iyer V, Offord V, Droop A, Behan F, Gonçalves E, Speak A, Iorio F, et al (2021) Combinatorial CRISPR screen identifies fitness effects of gene paralogues. Nat Commun 12: 1302

Tsherniak A, Vazquez F, Montgomery PG, Weir BA, Kryukov G, Cowley GS, Gill S, Harrington WF, Pantel S, Krill-Burger JM, et al (2017) Defining a Cancer Dependency Map. Cell 170: 564–576.e16

Wainberg M, Kamber RA, Balsubramani A, Meyers RM, Sinnott-Armstrong N, Hornburg D, Jiang L, Chan J, Jian R, Gu M, et al (2021) A genome-wide atlas of co-essential modules assigns function to uncharacterized genes. Nat Genet 53: 638–649

Williamson CT, Miller R, Pemberton HN, Jones SE, Campbell J, Konde A, Badham N, Rafiq R, Brough R, Gulati A, et al (2016) ATR inhibitors as a synthetic lethal therapy for tumours deficient in ARID1A. Nat Commun 7: 13837

Yates AD, Achuthan P, Akanni W, Allen J, Allen J, Alvarez-Jarreta J, Amode MR, Armean IM, Azov AG, Bennett R, et al (2019) Ensembl 2020. Nucleic Acids Res: gkz966

